# Natural variability increases human walking metabolic costs and its implications to simulation-based metabolic estimation

**DOI:** 10.1101/2025.03.13.643096

**Authors:** Aya Alwan, Manoj Srinivasan

**Affiliations:** Mechanical and Aerospace Engineering, The Ohio State University, 201, W. 19th Ave, Columbus, 43210, Ohio, United States

**Keywords:** stride-to-stride variability, metabolic cost, energetics, human walking, averaging, simulation, OpenSim, accuracy, Computed Muscle Control, Umberger energy model

## Abstract

Human walking contains variability due to small intrinsic perturbations arising from sensory or motor noise, or to promote motor learning. We hypothesize that such stride-to-stride variability may increase the metabolic cost of walking over and above a perfectly periodic motion, and that neglecting such variability in simulations may mis-estimate the metabolic cost. Here, we quantify the metabolic estimation errors accrued by neglecting the stride-to-stride variability using human data and a musculoskeletal model by comparing the cost of multiple strides of walking and the cost of a perfectly periodic stride with averaged kinematics and kinetics. We find that using an averaged stride underestimates the cost by about 2.5%, whereas using a random stride may mis-estimate the cost positively or negatively by up to 15%. As a further illustration of the cost increase in a simpler dynamical context, we use a feedback-controlled inverted pendulum walking model to show that increasing the sensory or motor noise increases the overall metabolic cost, as well as the variability of stride-to-stride metabolic costs, as seen with the musculoskeletal simulations. Our work establishes the importance of accounting for stride-to-stride variability when estimating metabolic costs from motion.

## 1 Introduction

Human locomotion does not follow a perfectly repetitive pattern and has deviations from nominal periodic motion trajectories that changes from stride-to-stride [1–5]. In the absence of obvious external perturbations or variability, for instance on a treadmill, this stride-to-stride variability could be due to small intrinsic perturbations arising from sensory or motor noise [6, 7], or potentially intentional variability used by the nervous system to promote motor learning [8–10]. In less constrained conditions, extrinsic factors such as changes in terrain and environmental conditions can contribute to the variability of walking patterns [11, 12]. To maintain stable walking motion despite such deviations and perturbations, humans use corrective feedback control strategies such as foot placement control [13–15] and ankle push-off modulation [13, 16]. These stabilizing control actions may potentially increase the overall muscle force and energetic demands. Here, using simulation models and experimental data on human walking, we examine how much such natural variability might contribute to increased metabolic cost.

While there is no definitive evidence on the causal effect of natural stride-to-stride variability on the total metabolic cost during human walking, increases in natural step length variations has been found to be linearly correlated with increases in metabolic cost measured through indirect calorimetry [17, 18] — although these experiments did not control for other covariates like speed, so it is not clear whether the increased cost is due to speed or variability. Step width variability induced by externally applied visual perturbations has been shown to increase measured metabolic cost [19], as do intentional step-length variability [18] or responses to other external perturbations [20]. Conversely, providing active stabilization and thereby reducing some stepping variability reduces metabolic cost [21] – although the springy stabilizing mechanisms in these experiments could have other energetically beneficial effects than just reducing variability. Taken together, such prior experiments lead us to the hypothesis that natural stride-to-stride variability may increase metabolic cost above the counter-factual of variability-free walking. Increased metabolic demand due to such variability may be attributed to (1) increase in muscle actuation resulting from sensory or motor noise leading to deviations from what is considered optimal, and (2) the muscle actuation and the energetic cost needed to correct these errors, so that the walking remains stable [4, 9, 14].

Understanding the metabolic implications of natural variability may potentially help improve estimations of walking metabolic costs. Estimating energy expenditure is useful as it allows us to quantify movement effort. Further, understanding the determinants of walking energetics is useful as there is substantial evidence that humans tend to move in a manner that minimizes their metabolic cost [22, 23]: this energy-optimality-based predictive theory has been used to explain many experimentally-observed aspects of legged locomotion [22–31]. To estimate metabolic costs from motion and ground reaction force data, researchers sometimes use musculoskeletal simulation software such as OpenSim [32, 33] in concert with metabolic models such as those to Umberger and Bhargava [34–36]. To perform such estimations of metabolic cost, researchers use motion and force data from a representative gait cycle or averaged data from multiple gait cycles, creating an averaged gait cycle, to estimate a single average metabolic cost [37–42] — potentially ignoring the stride-to-stride variability. While these simulation methods can predict or estimate large differences and broad trends in metabolic cost, they may not usually capture subtle changes or be quantitatively accurate [30, 43]. Hence, improving estimations of metabolic energy rates can improve our understanding of human movement and the development of interventions.

In this manuscript, we quantify the metabolic estimation errors accrued by neglecting the stride-to-stride variability, and propose a natural variant of traditional methods to better account for such affects. Specifically, we will estimate the total muscle metabolic cost of walking from experimental motion and force data using three methods (Figure 1): 1) the traditional method of using joint angles and ground reaction forces for a single stride (i.e. gait cycle) and then estimating a modeled metabolic cost, 2) the method of averaging joint angles and ground reaction forces for multiple strides and estimating a single modeled metabolic cost, and 3) the proposed method by estimating metabolic cost for multiple strides separately and then estimating one averaged metabolic cost value across these strides, with the goal of fully capturing the stride-to-stride variability. We apply these methods to metabolic estimations using the Umberger metabolic cost model [34–36, 44] and using a simple torque-squared model of effort [29, 45] commonly used in force distribution or static optimization applications [46–48], both using an OpenSim model [32] and long time-series human walking data [49]. As a complement to such use of detailed human data and musculoskeletal model, we also illustrate similar metabolic trends with a simple inverted pendulum walker [9, 28, 50–52], characterizing how sensory and motor noise increase stride-to-stride variability and the overall metabolic cost. We hypothesize that step-to-step variability contributes to greater effort, and hence, researchers may need to account for such cost addition due to intrinsic human perturbations.

**Fig. 1.**
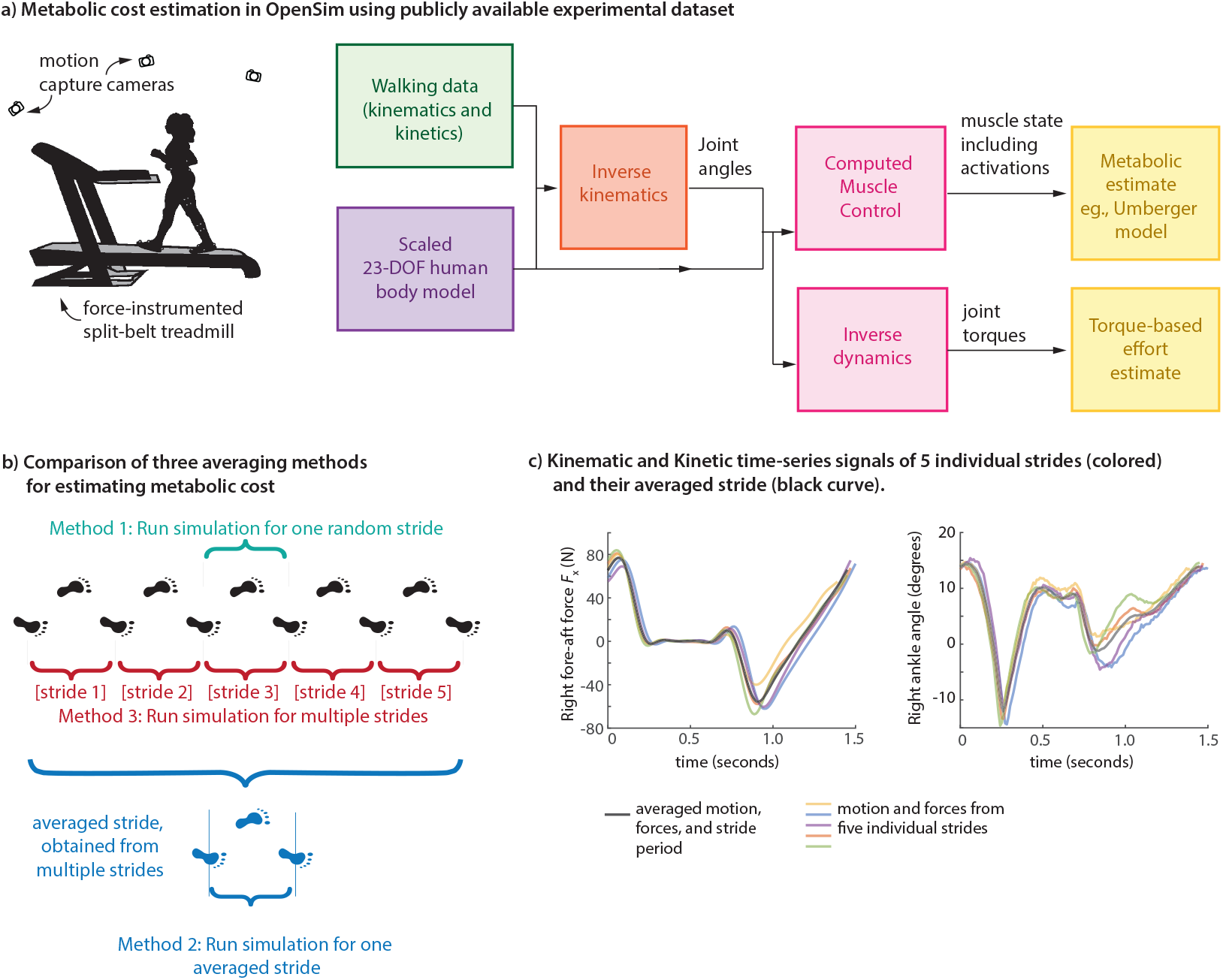
a) Metabolic cost estimation in OpenSim. Using marker motion and ground reaction force information from a multi-participant, multi-speed treadmill walking dataset, we obtain a metabolic cost estimate for each trial using a standard OpenSim process: using the Umberger metabolic model via computed muscle control for time-varying muscle state and using a joint-torque-based effort model via inverse dynamics for time-varying joint torques. b) Three averaging methods for estimating metabolic costs: estimating the cost for a randomly chosen individual stride (method 1), estimating the cost for the averaged gait pattern — averaged motion and ground reaction forces (method 2), and estimating an average cost over multiple strides (method 3). c) Obtaining an averaged gait pattern. Kinematic and kinetic time-series signals of five individual strides (colored) and their averaged version (black curve). Left panel shows the right foot fore-aft ground reaction force and the right panel shows the right ankle angle over one stride period of walking at 0.8 m/s.

## 2 Methods

### 2.1 Cost of variability from human walking data via a musculoskeletal model

#### Three metabolic cost averaging methods

We used OpenSim to estimate muscle energy expenditure based on publicly available experimental data [49] of human walking (Figure 1a). The dataset consists of walking trials for 15 subjects under normal and perturbed conditions. We used data from the first normal walking condition for 8 subjects [49]: 3 females and 5 males with an average age of 23.88 *±* 3.58 years, height of 1.759 *±* 0.061 meters, and mass of 72.66 *±* 13.51 kg. Treadmill data was used, so that multiple continuous strides of motion and force data was available. To estimate the effect of stride-to-stride variability on the metabolic cost, we used three different metabolic cost estimates: 1) metabolic cost calculated from a single randomly-selected stride (Figure 1b), 2) metabolic cost calculated from a single stride where the joint angles and ground reaction forces (GRFs) were averaged from *N*_stride_ strides (Figure 1b-1c), and 3) average metabolic cost calculated from *N*_stride_ individually simulated strides (Figure 1b). To obtain averaged motion and forces from multiple strides, we first mapped the data for each stride as going from 0% to 100% of the gait cycle, averaging the motion and force data at each gait cycle fraction to obtain an averaged function over a full gait cycle, and then mapping the resulting averaged motion or force pattern to the average stride period (e.g., Figure 1c). We treat approach 3 as being the proposed gold standard, taking into account the stride-to-stride variability fully, whereas approach 2 of using averaged motion and force patterns is equivalent to zeroing out the variability. The difference between these two approaches serves as an estimate of the metabolic penalty due to stride-to-stride variability. We used *N*_stride_ = 5 strides per walking trial for all the results reported here.

#### Metabolic cost modeling in OpenSim using the Umberger model

OpenSim [32, 33] provides tools to estimate muscle energy expenditure through its metabolic probes (Fig. 1a). Simulations of human walking were performed on the gait2354 musculoskeletal model, a lower extremity model with two legs and a lumped torso segment, with 23 degrees of freedom and 54 muscle-tendon actuators [32, 53]. We scaled the model to match the anthropometric properties of individual participants. We performed inverse kinematics using experimental motion data [49] to estimate joint angle trajectories [32, 33]. We used the inverse kinematics results from the residual reduction algorithm (RRA) tool as input to the Computed Muscle Control (CMC) calculation [32, 54]. The CMC calculation determines the muscle activations by using a feedback controller to have the resulting forward dynamic simulation track the experimentally observed kinematics and ground reaction forces [54]. The resulting muscle activations and dynamics are used to estimate time-varying metabolic rates via the Umberger metabolic cost model [34, 44, 55], with the metabolic cost integration and averaging performed in MATLAB (Fig. 1a).

The Umberger model available in OpenSim is a modified version of the muscle energetics model originally proposed by Umberger [34, 35, 44, 55], which calculates the total energy rate as a sum of work rate, 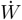, activation heat rate 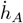, maintenance heat rate 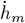, and muscle shortening and lengthening heat rate 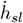 [34, 35, 37]. The estimated metabolic rate at any moment is a function of all the muscle state variables, namely, muscle activations, lengths, and shortening rates (which in turn also determine the force via a Hill model and mechanical power via standard physics).

#### Metabolic cost modeling using joint torques via inverse dynamics

As a simple alternative to the Umberger metabolic model, we also evaluated the effect of stride-to-stride variability on a joint-torque-based estimate of effort (Fig. 1a). Specifically, we defined the simple measure of effort *Ė*cost to be squared joint torques, summed over all joints and averaged over time:

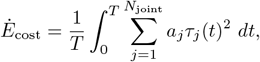

where *τ*_*j*_(*t*) is the moment at joint *j* at time *t, N*_joint_ is the total number of joints, *T* is the time duration over which the effort estimate is averaged, *a*_*j*_ is the joint-specific cost coefficient, assumed to be unity for the illustrative purposes here. The joint torques were obtained in OpenSim with the same human body model and walking experimental data as before.

#### Open gait data preparation and filtering

Publicly available open gait data by Moore et al. [49] were used in the analysis. We processed motion and force data from *N*_participant_ = 8 participants, each completing three trials of normal walking at three speeds, namely, 0.8 m/s, 1 m/s, and 1.2 m/s, for two minutes. Both motion and ground reaction force data were sampled at 100 Hz, with the motion capture data available for 19 markers being sufficient for scaling the OpenSim model and performing inverse kinematics. We recast the data in the OpenSim coordinate system (X is anterior-posterior, Y is vertical, and Z is medio-lateral) and processed both the inverse kinematics results and ground reaction force data with a low pass Butterworth filter with 6 Hz cutoff frequency. In OpenSim, for inverse dynamics, the ground reaction forces and moments were applied at the center of pressure (CoP): this meant that the horizontal ground reaction moments were zero by definition. Because the CoP is not defined when a foot is not on the ground (defined as when vertical force is less than 30 N), we replaced the unreliable CoP data during this period with a smooth interpolation of CoP from when the foot is on the ground: the details of the assigned CoP during this swing phase period do not affect the kinetics calculations such as inverse dynamics or CMC because the corresponding ground reaction forces are essentially zero. Left heel strikes were identified within each trial using 30 N vertical force threshold, and from these, six consecutive left heel strikes were randomly selected to produce five consecutive strides for further analysis within OpenSim as described in the previous paragraphs, specifically, computed muscle control and inverse dynamics.

### 2.2 Inverted pendulum model to illustrate the cost increase

As a complement to using 3D human gait data and musculoskeletal models, we use an inverted pendulum walking model [28, 50–52] to test the hypothesis that kinematic variability due to sensory or motor noise can increase the metabolic cost. The model involves walking stances phases that are inverted pendula, with step-to-step transitions that involve impulsive push-offs and heel-strikes [28, 50, 51]. The metabolic energy cost is modeled as due to the work done by the push-off impulse and an empirically based leg swing cost [56]. The nominal gait of this inverted pendulum walker is chosen to be energy optimal at a given speed [9]. Deviations from this nominal gait are corrected by an empirically based feedback controller that adjusts foot placement (step length) and push-off, depending on the forward speed during stance [1, 14]. Thus, the sensory input to the feedback controller is the forward speed and sensory noise is implemented as Gaussian noise added to this forward speed, resulting in stepping variability. Similarly, motor noise is modeled as Gaussian noise added to the push-off impulse and foot placement. The inverted pendulum model implementation in MATLAB and its parameters are identical to that used in a recent locomotor adaptation study [9]; the zero adaptation case used in this current manuscript with non-zero sensory and motor noise is available as Supplementary Material.

## 3 Results

### Umberger costs: costs have stride-to-stride variability and averaged gait pattern has lower costs

Within each walking trial, participants exhibited substantial stride-to-stride variability in the metabolic costs estimated using the Umberger model on individual strides (Figure 3a). Compared to the proposed gold standard of multiple-stride average of the estimated costs, the absolute percent error in using the cost of a randomly chosen individual stride ranged from about 0.03% to 7% at the walking speed of 0.8 m/s, 0.001% to 16% at 1.2 m/s, and 0.07% to 12% at 1.6 m/s, across all participants (Figure 3a). This spread of the percent error in individual stride metabolic estimates illustrate the stride-to-stride variability of the metabolic cost and evaluating the metabolic cost using a random stride may result in errors as large as 16% and this may be an over- or under-estimation.

Across all participants, the metabolic cost estimate using the averaged motion and force patterns resulted in lower average costs compared to averaging the costs across multiple strides: 2.3 *±* 0.67% (mean *±* s.e.) lower for 0.8 m/s, 3.3 *±* 0.3% lower for 1.0 m/s, and 1.9 *±* 0.6% lower for 1.2 m/s (Figure 3b). We consider this difference to be an estimate of the metabolic cost of stride-to-stride variability, about 2.5 *±* 0.07% across all participants and trials. Performing a paired t-test to determine whether the metabolic cost for the averaged gait pattern (method 2) differed significantly from the average costs over multiple strides (method 3) showed statistically significant differences across all speeds: *p* = 0.009 at speed 0.8 m/s, *p* = 0.018 at speed 1.2 m/s, and *p* = 0.002 at speed 1.6 m/s. Indeed, the metabolic cost of the averaged gait pattern was lower for 22 out of 24 trials across the three speeds and eight participants with the other 2 trials being negligibly higher (Figure 3a-b).

### Torque-based costs: costs have stride-to-stride variability and averaged gait pattern has lower costs

The metabolic trends for the simple torque-based effort cost was identical to the Umberger model costs. Within each walking trial, participants exhibited substantial stride-to-stride variability in the effort costs estimated using the torque-squared model on individual strides (Figure 3a). Compared to the gold standard of multiple-stride average of the estimated costs, the absolute percent error in using the cost of a randomly chosen individual stride ranged from about 0.003% to 14.6% at speed 0.8 m/s, 0.08% to 11% at speed 1.2 m/s, and 0.03% to 14.5% at speed 1.6 m/s across all participants (Figure 3c). Thus, choosing to estimate this torque-based effort using a random individual stride might result in an error as high as about 15%.

Across all participants, the torque-based effort cost estimate using the averaged motion and force patterns resulted in lower average costs compared to averaging the costs across multiple strides: 3.5 *±* 0.3% lower for 0.8 m/s, 4.9 *±* 1.1% lower for 1.0 m/s, and 4.1 *±* 0.9% lower for 1.2 m/s (Figure 3d). We consider this difference to be the metabolic cost of stride-to-stride variability, we estimate this cost to be about 4.2 *±* 0.1% lower when averaged across all participants and trials. Performing a paired t-test to determine whether the metabolic cost of the averaged gait pattern (method 2) differed significantly from the mean metabolic cost over multiple strides (method 3) showed statistically significant differences at all speeds: *p* = 0.005 at speed 0.8 m/s, *p* = 0.000 at speed 1.2 m/s, and *p* = 0.000 at speed 1.6 m/s. Indeed, the torque-based cost of the averaged gait pattern was lower than every one of the 24 trials across the three speeds and eight participants (Figure 3c).

### Insignificant speed dependence of metabolic estimation errors

We did not find a meaningful speed dependence of the metabolic errors. We fit a linear regression model with the trial speeds as the independent variables and percent error in metabolic cost as the dependent variable for the individual stride metabolic costs and the averaged gait pattern metabolic costs, for both the Umberger and the torque-based models. We did not find significant dependence of error on speed in either averaging methods for the torque-based model. Similarly, no speed-dependence trend was found for the averaged gait pattern-based cost for the Umberger model. However, there was a statistically significant (*p* = 0.037) increase in individual stride error as speed increases, but this dependence was very weak (about 1.5% error increase for a large 1 m/s speed increase).

### Inverted pendulum model illustrates cost of variability

The feedback-controlled inverted pendulum model (Figure 2a) exhibits kinematic stride-to-stride variability when subject to sensory noise or motor noise. This variability has a metabolic cost penalty, wherein the metabolic cost increases with the motor noise level (Figure 2b) and the sensory noise level (Figure 2c), compared to the no-noise baseline. Concomitant with this cost increase, the standard deviation of the stride-wise metabolic costs also increases with noise.

**Fig. 2.**
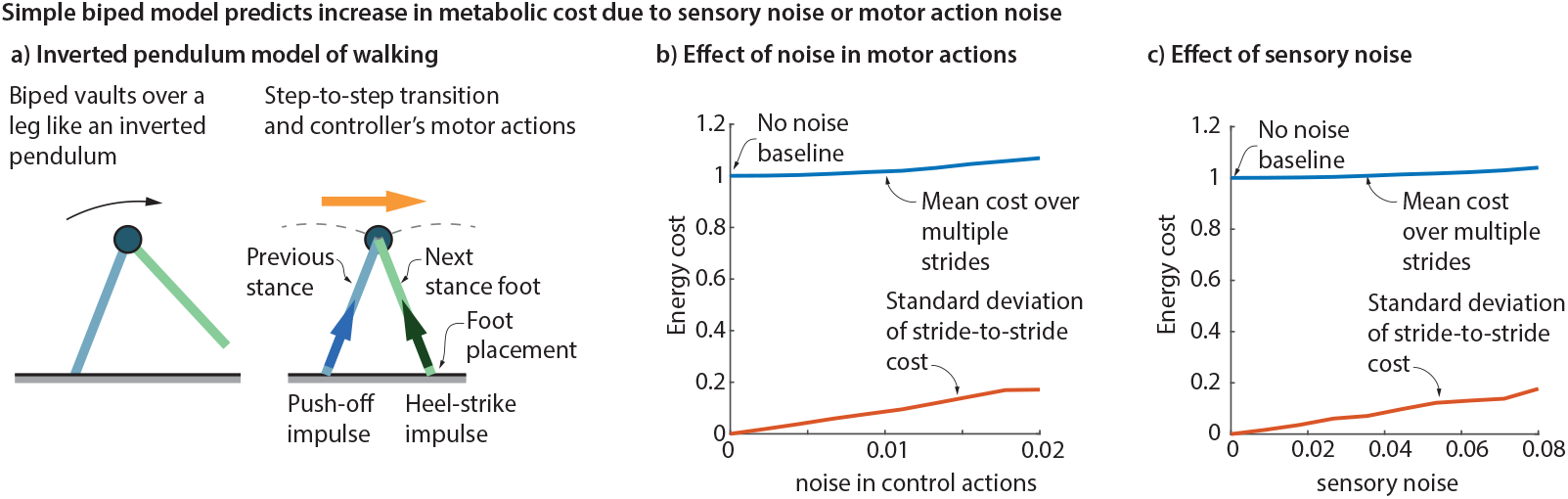
Cost of variability from an inverted pendulum walker. a) Inverted pendulum walking involves a compass gait during stance and a step-to-step transition that involves a push-off and a heel-strike impulse. b) Adding motor noise to push-off impulse and foot placement results in increased cost compared to the no-noise baseline. The standard deviation of the stride-to-stride cost also increases with the motor noise. c) Adding sensory noise to the stance phase speed, used as a sensory input to the walker’s feedback controller, results in higher cost compared to the no-noise baseline. The standard deviation of the stride-to-stride cost also increases with the sensory noise.

**Fig. 3.**
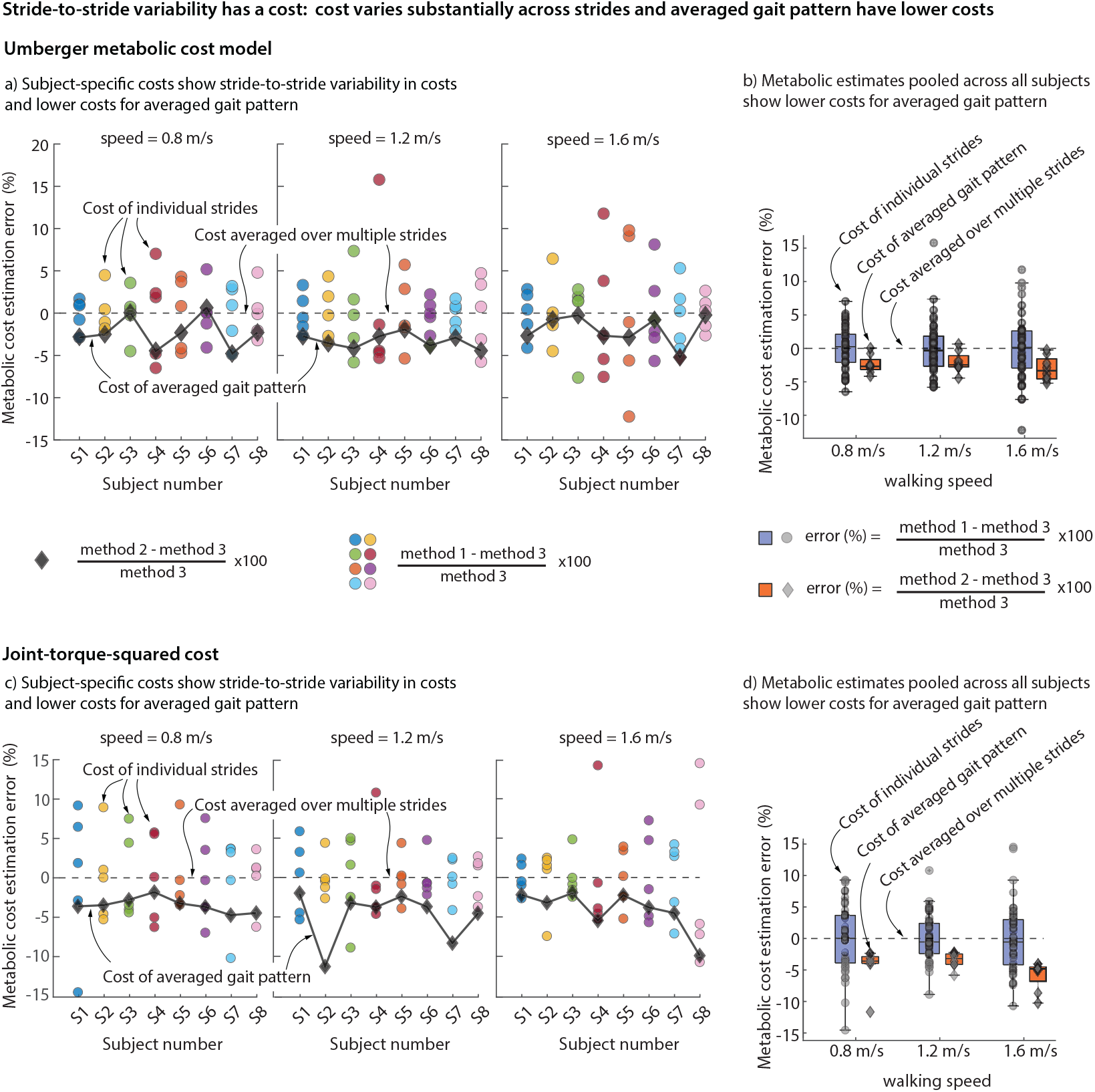
OpenSim simulation: Error in simulated metabolic cost between method 1) individual strides (circular scatter points) and method 2) averaged strides (diamond scatter points) from their method 3) individual stride arithmetic average, for eight participants walking at three speeds. Torque-based model: Error in simulated metabolic cost between method 1) individual strides (circular scatter points) and method 2) averaged strides (diamond scatter points) from their method 3) individual stride arithmetic average for eight participants walking at three speeds.

## 4 Discussion

We have demonstrated that there is substantial stride-to-stride variability in estimated metabolic or effort costs, so that using a randomly chosen individual stride to estimate the metabolic cost can produce errors of up to 15%, and this may be an over- or underestimation due to random variation. Further, by comparing a gold standard multi-stride estimate of metabolic cost with the cost of an averaged gait pattern – essentially with no stride-to-stride motion or force variation – we find that the metabolic penalty due to the stride-to-stride kinematic and kinetic variability is about 2.5% across multiple speeds, with the averaged gait pattern having a systematically lower cost and thus an underestimation of the true cost. Thus, the metabolic cost of stride-to-stride variability is small but significant. We found no significant speed dependencies of these errors. Our results suggest the importance of accounting for stride-to-stride variability influencing metabolic estimations. These insights not only advance our understanding of metabolic cost modeling, but also provide a foundation for refining predictive methods in human energetics within biomechanics.

The metabolic cost of stride-to-stride variability in walking may be due to some mixture of: (1) muscle actuation increase due to sensory noise, motor noise, and environmental variability and (2) the additional metabolic cost to correct the kinematic errors due to such noise, so that the walking remains stable. Because metabolic cost dependence on muscle activation and other muscle states may be nonlinear, the total metabolic penalty will not be additively separable into these component costs, even if the probability distributions of the noise and environmental variability were to be fully characterized and the stabilizing controller were to be fully understood.

The observation that the averaged gait pattern has systematically lower metabolic cost compared to the multi-stride average of the metabolic cost can be potentially understood via the Jensen’s inequality. Jensen’s inequality [57–59], in its simplest form (figure 4), states that for a convex function *f* (*p*), the function evaluated at the average value of two points *p*_1_ and *p*_2_, namely, *f* (*p*_1_ + *p*_2_)*/*2, is less than or equal to the average value of the function evaluated at those two points (*f* (*p*_1_) + *f* (*p*_2_))*/*2, i.e.,

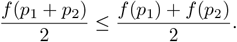

**Fig. 4.**
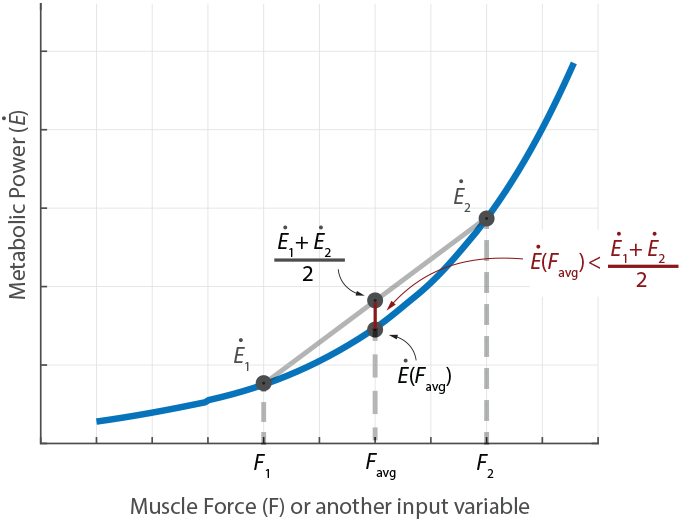
Metabolic power *Ė* as a function of muscle force *F* used to illustrate Jensen’s inequality.

More generally, if the variable *p* was time-varying *p*(*t*), again, the function evaluated at the average value of *p*(*t*) is less than or equal to the time-average of the time-varying function *f* (*p*(*t*)). For our purposes, we can identify the function *f* with the metabolic cost *Ė* and the variable *p* with muscle force or muscle activation or other relevant muscle state variables (see figure 4). The Umberger model is usually strictly convex in key input variables like muscle activation and muscle force, especially in the non-isometric regime (cost scales like activation squared) and linear in other variables like shortening velocity, so overall, we expect Jensen’s inequality to capture the underestimation. Other recent metabolic measurements find that metabolic cost scales faster than linearly even close to the isometric regime [45]. Of course, convexity of the true metabolic cost function is not established, so this is a model-based observation. Interestingly, the Umberger metabolic estimate for the averaged gait pattern was slightly higher for two trials compared to the multi-stride average cost: this does not necessarily violate the reasoning above, because our averages are over only five strides and the discrepancy may be due to, for instance, the participant’s gait being far from periodic and systematically slowing down, making reasoning based on Jensen’s inequality non-trivial. No such reversal of expectation was found for the simple torque-based cost. As an alternative to using Jensen’s inequality to explain the metabolic underestimate, we might posit that the energy optimal gait is perfectly stride-periodic and any deviation from it that is *N* -stride periodic would be non-optimal by definition and thus have higher energy. However, this reasoning is not a complete explanation of our observed results either, as it requires the observed motion to be optimal with respect to the specific cost functions used. But as we know, most predictive gait optimizations with different cost functions find non-trivial differences between the predicted motion and observed human data.

Estimating a model-based metabolic cost estimate as we have done here involves sources of error, including the use of simplified muscle and metabolic models, including neglecting potential force-rate costs [60], sensitivity to muscle-tendon parameter variations, inter-participant variations, and assumptions such as modeling muscles as massless. For computational tractability, we used five strides to estimate the more accurate metabolic cost and we expect the variance of the metabolic estimates to reduce further with more steps like 1*/N*_strides_ [61]. Our study does not include a validation using indirect calorimetry to estimate metabolic cost, as our goal was to estimate the cost of natural variability rather than variability that may be to external manipulations such as mechanical or sensory perturbations. While this may be considered a limitation, our objective was not necessarily to estimate an accurate metabolic cost for a particular application. Instead, we have provided a qualitative account of the potential errors in including or ignoring the effects of stride-to-stride variability and a rough model-based estimate of the cost of stride-to-stride variability.

We have estimated the errors due to ignoring stride-to-stride variability using treadmill data. Overground walking, even at steady state, is known to have greater speed fluctuations, and thus may have higher metabolic cost due to stride-to-stride variability [62]. While multiple continuous strides of both motion capture and ground reaction forces in overground walking may be challenging to obtain as most labs have a small number of force-plates, the multiple-strides analysis used herein are applicable as long as there are multiple strides of data, even if not continuous. Recently, researchers have proposed the use of machine-learned models of ambulatory locomotor metabolic costs with IMU-based sensors [63] and such models, as long as they are applied on multiple strides and averaged (and not computed on averaged strides or a random stride), are likely to account for the cost due to stride-to-stride variability, analogous to the results obtained herein. Future work could involve repeating this study in participants with movement disorders that may increase stride-to-stride variability [64, 65], thereby estimating the contribution of variability to increased metabolic cost.

In conclusion, we suggest that any future methods for metabolic cost estimation from motion and simulation should account for the effect stride-to-stride variability, so as to avoid unnecessary inaccuracies, especially in applications that involve subtle changes in metabolic costs.

## Acknowledgments

The work was supported in part by NIH grant R01GM135923-01, NSF SCH grant 2014506, and NSF CMMI grant 1254842.

## Competing interests

The authors declare that they have no competing interests.

## Notes

### Competing Interest Statement

The authors have declared no competing interest.

